# Infrared thermography cannot be used to approximate core body temperature in wild primates

**DOI:** 10.1101/2020.09.10.289512

**Authors:** Richard McFarland, Louise Barrett, Andrea Fuller, Robyn S Hetem, Warren Porter, Christopher Young, S Peter Henzi

## Abstract

Understanding the physiological processes that underpin primate performance is key if we are to assess how a primate might respond when navigating new and changing environments. Given the connection between an animal’s ability to thermoregulate and the changing demands of its thermal environment, increasing attention is being devoted to the study of thermoregulatory processes as a means to assess primate performance. Infrared thermography can be used to record the body surface temperatures of free-ranging animals. However, some uncertainty remains as to how these measurements can be used to approximate core body temperature. Here, we use data collected from wild vervet monkeys (*Chlorocebus pygerythrus*) to examine the relationship between infrared body surface, core body, and local climate, to determine to what extent surface temperatures reflect core body temperature. While we report a positive association between surface and core body temperature – a finding that has previously been used to justify the use of surface temperature measurements as a proxy for core temperature regulation – when we controlled for the effect of the local climate in our analyses, this relationship was no longer observed. That is, body surface temperatures were solely predicted by local climate, and not core body temperatures, suggesting that surface temperatures tell us more about the environment a primate is in, and less about the thermal status of its body core in that environment. Despite the advantages of a non-invasive means to detect and record animal temperatures, infrared thermography alone cannot be used to approximate core body temperature in wild primates.

## INTRODUCTION

As primate populations continue to decline and face an increasing risk of extinction as a consequence of climate change (Graham, Matthews, & Turner, 2016; Estrada et al., 2017), understanding the physiological processes that underlie the relationship between animal performance and environmental challenges is becoming increasingly important. Local climate exerts a strong selective pressure on animal behavior, physiology, and survivorship, and therefore has a profound impact on animal distributions and phenotypes (Hetem, Fuller, Maloney, & Mitchell, 2014; Fuller, Mitchell, Maloney, & Hetem, 2016; Mitchell et al., 2018). Given the obvious connection between an animal’s ability to thermoregulate and the demands of its thermal environment, increasing attention is being devoted to the study of body temperature as a means to assess primate performance in variable environments.

Early studies investigating primate body temperature regulation in response to environmental variability relied on measurements obtained from captive or laboratory-housed subjects (Sulzman, Fuller, & Moore-Ede, 1977; Whittow, Scammell, Manuel, Rand, & Leong, 1977; Wurster, Murrish, & Sulzman, 1985; Müller, Nieschalk, & Meier, 1985; McNab & Wright, 1987; Lubach, Kittrell, & Coe, 1992; Robinson & Fuller, 1999; Maloney, Mitchell, Mitchell, & Fuller, 2007). While this approach provides a high degree of experimental control, such studies do not provide the data to allow conclusions to be drawn about how a free-living animal responds to change in its natural habitat. Moreover, body temperatures in laboratory conditions typically are measured with thermosensitive probes or via telemetry with a receiver in close proximity, methods that are not readily transposable to studying primates in situ.

Advancements in remote telemetry or biologging have provided insights into the body temperature of a small number of free-ranging primate populations. Brain and Mitchell (1999) used intra-abdominal temperature-sensitive radio transmitters to record the core body temperature patterns of chacma baboons (*Papio ursinus)* under free-ranging, natural conditions. Their study was limited by relatively few subjects and body temperature measurements being collected only intermittently during the daytime over less than a month for each animal. Nevertheless, this study revealed the importance of high heat load as a thermal stressor to chacma baboons, and the importance of drinking water to prevent hyperthermia (Brain & Mitchell, 1999). Intra-abdominal transmitters also have been used to describe the core body temperature patterns associated with daily torpor in lemurs (*Lemuridae spp*. Schmid 2000; Schmid, Ruf, & Heldmaier, 2000; Dausmann, 2005). More recently, we have used intra-abdominal data loggers to describe the seasonal patterns of core body temperature in wild vervet monkeys (*Chlorocebus pygerythrus*), the thermoregulatory consequences of both cold and heat stress, inter-individual differences in thermal performance, and the importance of behavioral (including social) thermoregulation in body temperature regulation (Lubbe et al., 2014; McFarland et al., 2015; Henzi et al., 2017; McFarland, Henzi, & Barrett, 2019; McFarland et al., 2020).

The use of intra-abdominal data loggers provides a good index of the thermal status of the body core, but requires animals to be captured and to undergo surgical procedures for logger implantation and extraction (see McFarland et al., 2015 for a full description of methods). As an alternative to data logging approaches there has been interest in using infrared thermography to make inferences about thermoregulatory processes in free-ranging animals, including primates (McCafferty, 2007; Cilulko, Janiszewski, Bogdaszewski, & Szczygielska, 2013; Thompson et al., 2017; Narayan, Perakis, & Meikle, 2019). At face value, this appears an attractive option for non-invasive and remote measurement of body temperature. However, infrared thermography measures animal surface temperatures, and there is contrasting evidence whether surface temperatures (i.e., skin, fur, inner-ears, eyes) can approximate core body temperature. While some studies report no significant association between body surface and core body temperatures (Jay et al., 2007; Larcombe, 2007; Sikoski et al., 2007; Sykes et al., 2012), others have shown a positive correlation (Dausmann, 2005; Warriss, Pope, Brown, Wilkins, & Knowles, 2006; Johnson, Rao, Hussey, Morley, & Traub-Dargatz, 2011; Giloh, Shinder, & Yahav, 2012; Zanghi, 2016). An important caveat, however, is the lack of control in these analyses for the mediating effect that local climate has on both surface and core body temperatures. That is, even though body surface and core body temperatures were shown to be positively correlated, it is possible that this relationship was an artefact of the local climate influencing these variables. While it may be reasonable to assume that skin temperature more closely reflects core body temperature in small animals, the mediating effect of local climate should still be considered when attempting to make inferences about core body temperature from surface temperatures alone. In an experimental study of bats (*Carollia perspicillata*) in controlled environmental conditions, for example, it was observed that the difference between an animal’s core and skin temperature was a function of ambient temperature, with the authors concluding that any inferences made about core body temperature regulation from skin temperature measurements, including in very small animals, should account for the effect of ambient conditions (Audet & Thomas, 1996). Even when care has been taken to avoid increased surface heat loads incurred by solar radiation (McCafferty, 2007; Thompson et al., 2017), surface temperatures are still more influenced by local climate than body temperature. In wild mantled howling monkeys (*Alouatta palliata*), for example, dorsal fur temperature was more strongly predicted by ambient temperatures than by subcutaneous temperatures, and facial skin surface temperatures were predicted solely by ambient temperature and not by subcutaneous temperatures (Thompson et al., 2017).

Here, we use data collected from wild vervet monkeys to examine the relationship between body surface and core body temperatures, while controlling for local climatic conditions. Because an endothermic animal’s core body temperature is typically buffered from their environment by a range of autonomic and behavioural processes (Lovegrove, Heldmaier, & Ruf, 1991; Angilletta, Cooper, Schuler, & Boyles, 2010; Hetem, Maloney, Fuller, & Mitchell, 2016; Mitchell et al., 2018), we predict that surface temperatures will be more closely associated with local climate than with core body temperature. A better sense of the relationship between these variables will hopefully inform us whether infrared thermography can be used to approximate core body temperature. If surface temperatures are predicted by core temperatures, while controlling for the effect of the local climate, this would suggest that surface temperatures can approximate, to some extent, core body temperature. However, if surface temperatures are predicted by the local climate, and not core body temperatures, this would suggest that the surface temperatures tell us more about the environment an animal is in, and less about the body temperature of an animal in that environment. We use four body regions to test our hypothesis: the furred dorsal, ventral and tail surface and bare-skin facial surface.

## METHOD

In June 2017, as part of a longitudinal project on vervet monkey thermoregulation in the Eastern Cape, South Africa (32°22’S, 24°52’E), we collected infrared thermography data from a subset of individuals living across three groups (N=14: 5 females, 9 males). These animals fed on a natural diet, were fully habituated to the presence of researchers, and were individually identifiable by means of natural markings (Pasternak et al., 2013; McFarland, Barrett, Boner, Freeman, & Henzi, 2014).

### Infrared thermal imagery

We collected infrared thermal images (N_total_=294 images, 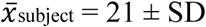 14 images) opportunistically using a handheld infrared thermograph model T360 camera (FLIR^®^ Systems Inc.). We set emissivity to 0.98. We targeted the animal’s furred dorsal, ventral, and tail surface, and bare-skin face surface, in each photo, to the point that some images could be used to measure multiple surface regions, and others not. We used the following selection criteria to determine the usability of infrared surface temperature measurements: (i) the animal was in plain view and not obscured by foliage or another individual, (ii) the animal was within 5 m of the camera, (iii) the targeted body surface was orientated toward the camera to avoid detection errors associated with sampling curved surfaces (McCafferty, 2007), and (iv) the sampled region of the body surface was in shade or low light levels, avoiding reflective light, thereby minimizing the effect of direct solar radiation on surface measurements (McCafferty, 2007). Table 1 outlines the distribution of the final measurements used in the current analyses.

**Table 1.** Sample of miniglobe, core body, and infrared body surface temperature measurements across four sites recorded from wild vervet monkeys

|  | Miniglobe | Core body | Body surface |  |  |  |
| --- | --- | --- | --- | --- | --- | --- |
|  |  |  | Dorsal | Ventral | Face | Tail |
| Number of monkeys | - | 14 | 14 | 11 | 13 | 14 |
| Number of images | 242 | 283 | 202 | 42 | 82 | 107 |
| Minimum temperature °C | 13.0 | 38.2 | 16.2 | 20.5 | 16.9 | 14.8 |
| Maximum temperature °C | 32.7 | 39.8 | 37.6 | 34.4 | 34.4 | 37.5 |
| Mean temperature ± SD °C | 23.6 ± 5.7 | 38.9 ± 0.3 | 26.3 ± 4.2 | 29.1 ± 3.75 | 27.0 ± 4.01 | 26.8 ± 4.9 |

At the time each image was taken, we positioned a thermocouple attached to a matte-black painted metal box within the frame of each thermal image (e.g., Figure 1). We calibrated the temperature measurements recorded by the thermal camera using the thermocouple. For each infrared image, we used FLIR Tools^®^ software (FLIR^®^ Systems Inc., 2019) to extract five discrete temperature measurements from the targeted body surface region and the black calibration box. We calculated the mean body surface and black box temperatures were for each image. We used the difference between the thermocouple temperature and the mean FLIR temperature of the black box to create a single-point calibration offset for all mean body surface temperatures.

**Figure 1.**
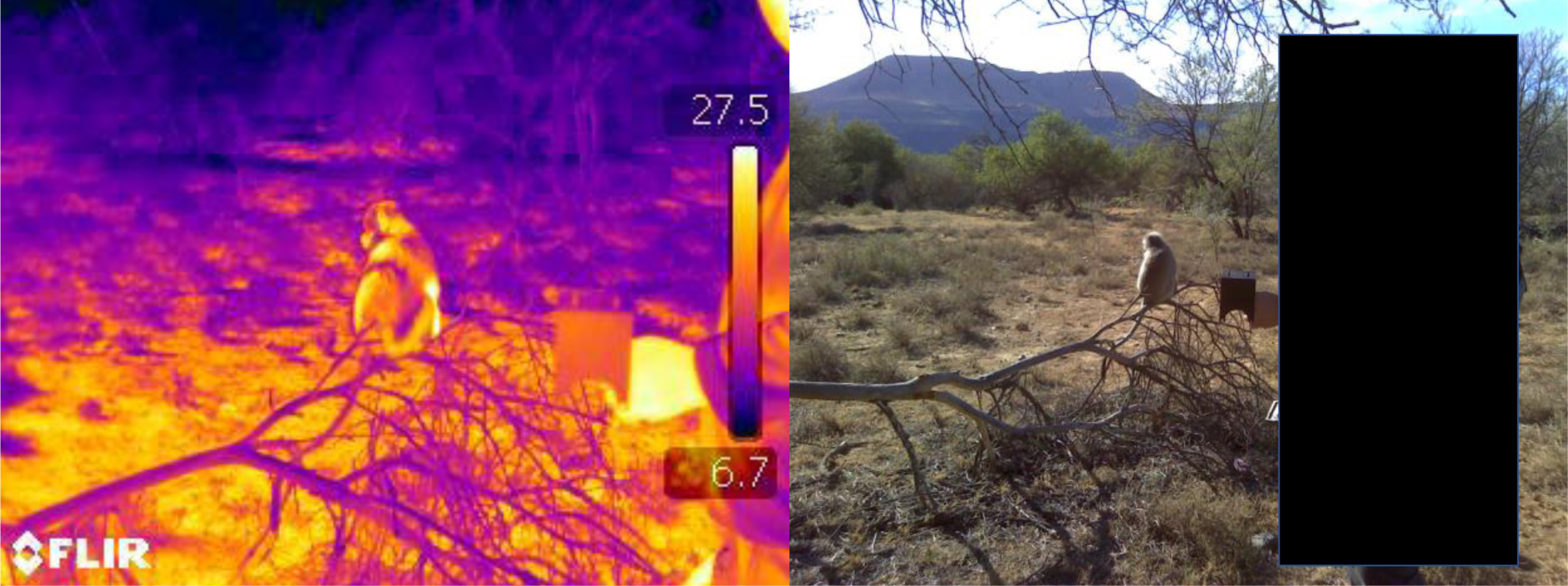
An infrared image and time-matched photograph of data collection (zoomed out for illustrative purposes) of a vervet monkey implanted with a temperature-sensitive data logger

### Local climate

We used black globe temperature to measure the local climate experienced by our study animals. Black globe temperature integrates the influence of air temperature, solar radiation and wind speed, and is thus considered a better measure of the thermal heat load experienced by an animal than air temperature alone (Hetem, Maloney, Fuller, Meyer, & Mitchell, 2007; McFarland et al., 2014). We recorded black globe temperatures every minute using a calibrated, temperature-sensitive Thermochron 4K iButton (resolution = 0.06°C; model DS1921G; Maxim Integrated™) housed inside a matte-black painted copper ball with a diameter of 30mm (hereafter, miniglobe). We recorded miniglobe temperatures at the time each image was taken, within 5m of the target animal. Miniglobe temperatures ranged from 13.0-32.7°C across the study period with a mean daily temperature of 23.6 ± SD 5.7°C.

### Body temperature data

In June 2016, we surgically implanted, intra-abdominally, 14 adult vervet monkeys (5 females, 9 males; distributed across three groups) with miniature temperature-sensitive data loggers (model: DST Centi-T, Star-Oddi, Iceland). Data loggers recorded core body temperature at five-minute intervals at a resolution of 0.03°C and were individually calibrated to an accuracy of 0.1°C. We recorded the body mass (kg) of all animals (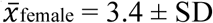0.4g, 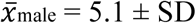 0.3g). We removed the data loggers at the end of June 2017. For full details of the capture and surgery procedure see McFarland et al. (2015). Data logger temperatures data were used to measure the core body temperature (to the nearest five minutes) of the same animals sampled using the infrared camera. Importantly, we implanted our subjects with data loggers in fulfilment of a long-term study of vervet monkey thermal physiology. That is, our subjects were not solely exposed to this procedure for the purpose of the current project.

Observational data collection protocols were approved by the University of Lethbridge under Animal Welfare Protocols 0702 and 1505. Capture and surgical procedures were approved by the University of the Witwatersrand Animal Ethics Research Committee (Protocol # 2015-04-14B-2017), and were treated in accordance with international ethical standards. No long-term sequelae were observed as a consequence of surgical intervention. Overall, this study adhered to the legal requirements of South Africa, and the American Society of Primatologists (ASP) Principles for the Ethical Treatment of Non-Human Primates.

### Statistical analysis

We used a series of Bland-Altman plots to visually compare the measurement of vervet monkeys’ core body temperature using intra-abdominal data loggers, with surface temperature measured using infrared thermography (Altman & Bland, 1983).

We performed our analyses in R v.3.6.0 (R-Core-Team, 2019) using the ‘lme4’ package to model outcomes (Bates et al., 2019), the ‘rsq’ package to generate adjusted *R*^*2*^ values for the fixed effects (Zhang, 2018), and the ‘lmertest’ package to generate p-values (Kuznetsova, Brockhoff, Christensen, & Jensen, 2019). Prior to running each model, we checked for intercollinearity by calculating variance inflation factors (VIF) for the predictor variables using the ‘car’ package (Fox et al., 2020). The VIF scores in all of our models were < 2 and were therefore not considered collinear. We scaled and centered our predictor variables so we could directly compare the resulting coefficients. We specified a gaussian error structure with a log link function in all of our models to normalize the residuals.

We originally ran a series of linear mixed models, entering subject ID as a random effect. However, these random effects did not explain any meaningful variance and their inclusion produced overfitted models with singular fit. We therefore ran a series of linear models, removing this random effect. The results of the linear mixed models and linear models were qualitatively the same. We present the results of the linear models below.

We first ran a series of four linear models entering our four surface temperatures (i.e., dorsal, ventral, tail, and face) in turn as the dependent variable, and time-matched miniglobe as the sole predictor variable. We ran a second series of four linear models entering our four surface temperatures in turn as the dependent variable, and time-matched core body temperature as the sole predictor variable. We ran a third series of four linear models entering our four surface temperatures in turn as the dependent variable. We entered time-matched miniglobe temperature and core body temperature as predictor variables, to determine whether this model improved upon either of the single predictor variable models described in the first two series of linear models.

For the first two series of linear models (i.e., the single predictor variable models), we follow Colquhoun (2014) in describing outcomes as indicating weak (P ∼ 0.05), moderate (P ∼ 0.01) or strong (P ∼ 0.001) evidence for effects. Following the third series of linear models, we used a reduction in the Akaike Information Criterion (AIC: Akaike, 1974) of a model, using a ΔAICc (to control for small sample sizes) threshold of < −2.0 (Burnham & Anderson, 2002), to indicate whether the composite model was better than its single predictor variable equivalent. We used adjusted *R*^*2*^ values to describe how much variance in a model’s dependent variable was explained by its predictor variables.

## RESULTS

In a series of four Bland-Altman plots (Figure 2), we observed large differences between the measurements of core body and surface temperatures 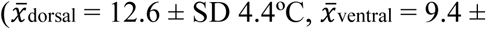 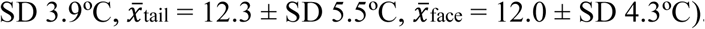. Temperature differences in excess of 15°C were common and occasionally exceeded 20°C at low temperatures. When the mean of core body and surface body temperatures were higher as a result of increased surface temperature in warm conditions (Table 2), the differences between these variables were smaller. However, core body and surface temperatures were never equivalent.

**Table 2.** Results of the linear model analyses testing the relationship between miniglobe temperature and infrared surface temperatures from the following body regions: dorsal, ventral, tail, and face. We ran the analyses at the level of the image/subject.

| | $\beta \pm SE$ | $t$ | $P$ |
| --- | --- | --- | --- |
| **Dorsal** (N=153 images, N=14 subjects) |  |  |  |
| Miniglobe temperature | 0.16 ± 0.01 | 25.02 | <0.001 |
| Intercept | 3.26 ± 0.01 | 532.10 | - |
| *Adjusted R<sub>2</sub>* (%) | 81.45 |  |  |
| *AICc* | 637.31 |  |  |
| **Ventral** (N=32 images, N=11 subjects) |  |  |  |
| Miniglobe temperature | 0.12 ± 0.01 | 9.30 | <0.001 |
| Intercept | 3.37 ± 0.01 | 274.70 | - |
| *Adjusted R<sub>2</sub>* (%) | 75.56 |  |  |
| *AICc* | 139.81 |  |  |
| **Tail** (N=76 images, N=14 subjects) |  |  |  |
| Miniglobe temperature | 0.21 ± 0.01 | 20.27 | <0.001 |
| Intercept | 3.26 ± 0.01 | 345.44 | - |
| *Adjusted R<sub>2</sub>* (%) | 86.75 |  |  |
| *AICc* | 328.56 |  |  |
| **Face** (N=63 images, N=13 subjects) |  |  |  |
| Miniglobe temperature | 0.14 ± 0.01 | 14.26 | <0.001 |
| Intercept | 3.28 ± 0.01 | 322.88 | - |
| *Adjusted R<sub>2</sub>* (%) | 76.69 |  |  |
| *AICc* | 276.80 |  |  |

**Figure 2.**
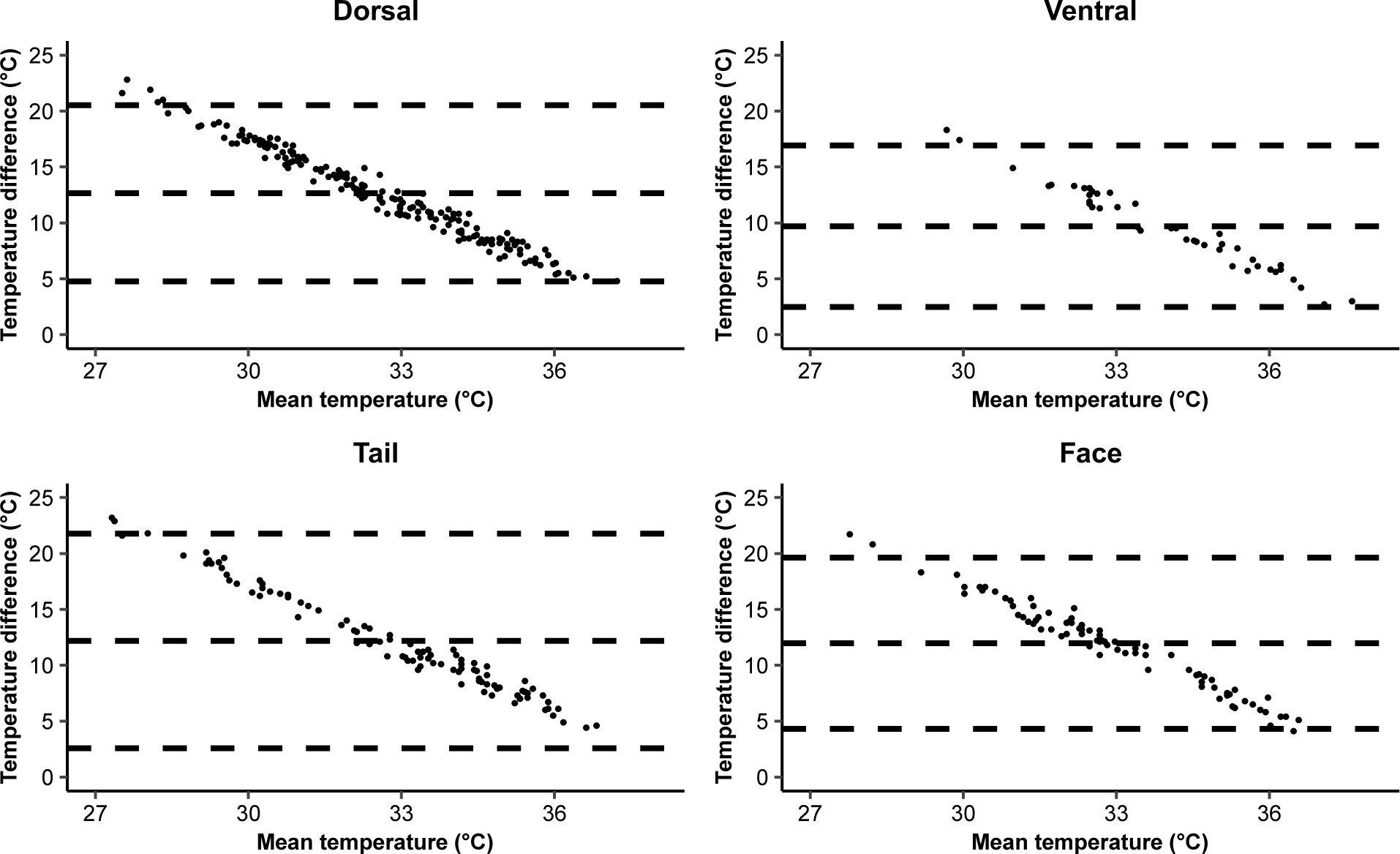
A Bland-Altman plot showing the relationship between vervet monkeys’ core body temperature (°C) and each of the four body surface temperatures (°C) (i.e., dorsal, ventral, tail and face). The x-axis represents the combined mean of core body and specified surface temperatures (°C). The y-axis represents the difference in temperature between core body and specified surface temperatures. Dashed lines represent the mean ± 1.96SD.

### Miniglobe temperature

In a series of four linear models (Table 2) with miniglobe temperature as the sole predictor variable, globe temperature had a strong positive effect on dorsal, ventral, face, and tail infrared surface temperatures (all P<0.001). Miniglobe temperature explained 81% (dorsal), 76% (ventral), 87% (tail), and 77% (face) and of the variance in infrared body surface temperatures (Table 2).

### Core body temperature

In a series of four linear models with core body temperature as the sole predictor variable, core body temperature had a strong positive effect on dorsal surface temperatures (P<0.001) and a weak positive effect on ventral (P=0.04), tail (P=0.02) and face (P=0.048) infrared surface temperatures. Core body temperature explained 8% (dorsal), 11% (ventral), 6% (tail), and 5% (face) of the variance in infrared body surface temperatures (Table 3).

**Table 3.** Results of the linear model analyses testing the relationship between **core body temperature** and infrared surface temperatures from the following body regions: dorsal, ventral, tail, and face. We ran the analyses at the level of the image/subject.

| | $\beta \pm SE$ | $t$ | $P$ |
| --- | --- | --- | --- |
| **Dorsal** (N=153 images, N=14 subjects) |  |  |  |
| Core body temperature | 0.05 ± 0.01 | 3.91 | <0.001 |
| Intercept | 3.27 ± 0.01 | 247.92 | - |
| *Adjusted R<sub>2</sub>* (%) | 8.24 |  |  |
| *AICc* | 881.92 |  |  |
| **Ventral** (N=32 images, N=11 subjects) |  |  |  |
| Core body temperature | 0.05 ± 0.02 | 2.18 | 0.04 |
| Intercept | 3.38 ± 0.02 | 146.63 | - |
| *Adjusted R<sub>2</sub>* (%) | 10.58 |  |  |
| *AICc* | 181.31 |  |  |
| **Tail** (N=76 images, N=14 subjects) |  |  |  |
| Core body temperature | 0.06 ± 0.02 | 2.39 | 0.02 |
| Intercept | 3.28 ± 0.02 | 138.46 | - |
| *Adjusted R<sub>2</sub>* (%) | 5.80 |  |  |
| *AICc* | 477.62 |  |  |
| **Face** (N=63 images, N=13 subjects) |  |  |  |
| Core body temperature | 0.04 ± 0.02 | 2.01 | 0.048 |
| Intercept | 3.29 ± 0.02 | 164.53 | - |
| *Adjusted R<sub>2</sub>* (%) | 4.64 |  |  |
| *AICc* | 365.57 |  |  |

### Miniglobe temperature and core body temperature

Given that globe temperature was a stronger predictor of infrared surface temperatures than core body temperature (Tables 2 and 3), we ran a series of four linear models including both globe temperature and core body temperature as predictor variables, to examine whether the addition of core body temperature improved the performance of the model with globe temperature as the sole predictor (Table 4). The dorsal, ventral, tail, and face infrared surface temperature models were not improved by the addition of core body temperature. All ΔAICc were > −2.0 with < 1% additional variance explained in all cases (Table 5). The best fitting models, for all infrared body surface temperatures, therefore, were those that only included globe temperature as a sole predictor variable.

**Table 4.** Results of the linear model analyses testing the effects of miniglobe temperature and core body temperature on infrared surface temperatures from the following body regions: dorsal, ventral, tail, and face. We ran the analyses at the level of the image/subject.

| | $\beta \pm SE$ | $t$ | $P$ |
| --- | --- | --- | --- |
| **Dorsal** (N=153 images, N=14 subjects) |  |  |  |
| Miniglobe temperature | 0.15 ± 0.01 | 23.76 | <0.001 |
| Core body temperature | 0.01 ± 0.01 | 1.82 | 0.07 |
| Intercept | 3.26 ± 0.01 | 536.18 | - |
| *Adjusted R<sub>2</sub>* (%) | 81.73 |  |  |
| *AICc* | 636.12 |  |  |
| **Ventral** (N = 32 images, N = 11 subjects) |  |  |  |
| Miniglobe temperature | 0.13 ± 0.15 | 8.34 | <0.001 |
| Core body temperature | -0.01 ± 0.01 | -0.46 | 0.65 |
| Intercept | 3.37 ± 0.01 | 270.99 | - |
| *Adjusted R<sub>2</sub>* (%) | 74.90 |  |  |
| *AICc* | 142.20 |  |  |
| **Tail** (N = 76 images, N = 14 subjects) |  |  |  |
| Miniglobe temperature | 0.21 ± 0.01 | 19.29 | <0.001 |
| Core body temperature | 0.00 ± 0.01 | 0.38 | 0.70 |
| Intercept | 3.26 ± 0.01 | 343.47 | - |
| *Adjusted R<sub>2</sub>* (%) | 86.59 |  |  |
| *AICc* | 330.64 |  |  |
| **Face** (N = 63 images, N = 13 subjects) |  |  |  |
| Miniglobe temperature | 0.14 ± 0.01 | 13.48 | <0.001 |
| Core body temperature | 0.00 ± 0.01 | 0.10 | 0.92 |
| Intercept | 3.28 ± 0.01 | 320.45 | - |
| *Adjusted R<sub>2</sub>* (%) | 76.31 |  |  |
| *AICc* | 279.08 |  |  |

**Table 5.** A summary of the performance (AICc and Adjusted R2 %) of the infrared surface temperature linear models (i.e., dorsal, ventral, face, and tail) as explained by miniglobe temperature and core body temperature as sole predictors, and a composite model including both predictor variables. ^†^ denotes best fitting model. A ΔAICc threshold of < −2.0 was used indicate whether a composite model was better than its single predictor variable equivalent.

|  |  | Dependent variables |  |  |  |
| --- | --- | --- | --- | --- | --- |
| Predictor variables | | Miniglobe temperature | Core body temperature | Miniglobe temperature *and* Core body temperature | $\Delta$ change in performance of the mini globe model with the addition of core body temperature |
| **Dorsal** | AICc | 637.31† | 881.92 | 636.12 | -1.19 |
|  | R<sub>2</sub> | 81.45 | 8.24 | 81.73 | 0.28 |
| **Ventral** | AICc | 139.81† | 181.92 | 142.2 | 2.39 |
|  | R<sub>2</sub> | 75.56 | 10.58 | 74.9 | -0.66 |
| **Tail** | AICc | 328.56† | 477.62 | 330.64 | 2.08 |
|  | R<sub>2</sub> | 86.75 | 5.8 | 86.59 | -0.16 |
| **Face** | AICc | 276.80† | 365.57 | 279.08 | 2.28 |
|  | R<sub>2</sub> | 76.69 | 4.64 | 76.31 | -0.38 |

## DISCUSSION

Our findings reveal that infrared thermography cannot be used to approximate core body temperature. Vervet monkey surface temperatures measured on the furred dorsal, ventral and tail region, as well as the bare-skin facial region, were more strongly predicted by miniglobe (i.e., environmental) temperatures than core body temperatures. At low ambient temperatures, surface temperatures dropped substantially and furred surfaces could be more than 20°C below core body temperature. Since an animal’s body surface is the interface where heat is gained by, and dissipated from, the body, it is unsurprising that animal surface temperature is strongly influenced by environmental temperature. While the core body temperatures of free-ranging primates can also vary in response to environmental variability (e.g., Brain & Mitchell, 1999; Schmid, 2000; Schmid et al., 2000; Dausmann, 2005; Lubbe et al., 2014; McFarland et al., 2015, Henzi et al., 2017; McFarland et al., 2020), core body temperature is much more tightly regulated through a range of behavioral and physiological processes.

Primates, like all mammals, employ a range of physiological mechanisms to cope with environmental challenges while maintaining homeostasis. Autonomic processes involve the activation of pathways in the preoptic area of the hypothalamus that regulate the balance of heat production and loss (Morrison & Nakamura, 2019). Individuals can also engage in behaviors that alter body temperature, such as changing activity patterns, posture, or selecting appropriate microclimates (McFarland et al., 2015, 2019, 2020; Henzi et al., 2017). Heat transfer to and from the body’s core is also buffered by the conductive and reflective properties of the pelt (i.e., color, depth, density, and condition: Scholander, 1950; Schmidt-Nielsen, 1997; McFarland et al., 2016). Given the complexities surrounding core body temperature regulation, it would be surprising if surface temperatures did approximate core body temperatures. Yet, a number of studies have shown an association between surface temperature and core body temperature (Dausmann, 2005; Warris et al., 2006; Johnson et al., 2011; Giloh et al., 2011; Zanghi, 2016), including the current manuscript. However, the variance in surface temperature that could be explained by core body temperature in our study was an order of magnitude lower than that explained by miniglobe temperature. In addition, after controlling for the effect of miniglobe temperature, the relationship with surface and core temperatures became trivial. We recommend that future studies do not use surface temperature measurements to approximate core body temperature, without a full understanding of the interactions between core, surface, and environmental variables.

That is not to say that infrared thermography cannot provide important insights on thermoregulatory processes. Surface temperatures not only inform us about the heat load experienced at an animal’s surface, but may also provide information on peripheral blood flow, and the effect of pelage properties on heat transfer (McCafferty, 2007; Cilulko et al., 2013; Mathewson et al., 2018). Infrared thermography can also provide valuable insights on animal behavior and physiology, or to test the accuracy of biophysical heat transfer models (e.g., Mathewson et al., 2018). In primates, infrared thermography has been used to assess the behavior of otherwise unobservable nocturnal species (e.g., Rahlfs & Fichtel, 2009; Tan, Yang, & Niu, 2013; Chen et al., 2015). Coupled with drone technology, infrared thermography has been used to provide population estimates by detecting animal heat-signatures in remote, difficult-to-navigate locations (Kays et al., 2018; Spaan et al., 2019; Burke et al., 2019). Facial infrared thermal imaging has been used to quantify the emotional states of non-human primates (Nakayama, Goto, Kuraoka, & Nakamura, 2005; Kuraoka & Nakamura, 2011; Ioannou, Chotard, & Davila-Ross, 2015; Chotard, Ioannou, & Davila-Ross, 2018) and other animals (McCafferty, 2007).

In an apparent attempt to avoid the invasiveness of intra-abdominal data logging, several research teams have used skin or subcutaneous temperature measurements to make inferences about core body temperature in primates. These methods have provided information on the hibernation patterns of the Lesser bushbaby (*Galago moholi*: Mzilikazi, Masters, & Lovegrove, 2006; Nowack, Mzilikazi, & Dausmann, 2010; Nowack, Wippich, Mzilikazi, & Dausmann, 2013) and several lemur species (*Lemuridae spp*., Schmid, 2001; Dausmann, Glos, Ganzhorn, & Heldmaier, 2004; Dausmann, 2005; Blanco, Dausmann, Faherty, & Yoder, 2013; Kobbe, Nowack, & Dausmann, 2014), as well as the seasonal variability in the body temperature rhythm of the larger, diurnal mantled howling monkey (Thompson et al., 2014). While subcutaneous body temperatures, typically recorded using devices implanted between the scapula, may more closely reflect core body temperature than surface temperature, they also are likely to be influenced by a core to periphery gradient, particularly in large mammals. The temperature of peripheral tissue is more strongly influenced by local climate, and peripheral blood flow, than is core body temperature (Mitchell et al., 2018). The subcutaneous body temperatures of mantled howling monkeys, for example, were strongly influenced by environmental temperature (Thompson et al., 2014). Furthermore, although a positive association between core body and subcutaneous body temperatures has been documented (Brown & Bernard, 1991; Navarro-Serra & Sanz-Cabañes, 2018), this does not mean that the measures are equivalent or interchangeable. Similar to surface temperatures, subcutaneous temperatures may deviate substantially from core body temperature, particularly in cold environments when endotherms peripherally vasoconstrict to conserve core body heat (Torrao, Hetem, Meyer, & Fick, 2011).

Other less invasive measures of an animal’s body temperature include measuring the temperature of an animal’s feces to approximate its core body temperature. For example, fecal temperature has been used as a proxy for core body temperature in chimpanzees (*Pan troglogytes*: Jensen, Mundry, Nunn, Boesch, & Leendertz, 2009; Negrey, Sandel, & Langergraber, 2020) based on limited evidence that fecal temperatures approximate rectal temperature in humans. However, given the infrequency of defecation point-sampling, and the fact that core body temperatures can fluctuate over 24h, and over even shorter time intervals for a multitude of reasons (e.g., drinking, behavioral thermoregulation, microclimate selection, intensity of activity: McFarland et. al., 2015, 2019, 2020), this method offers very little information on the regulation of core body temperature. Continuous and remote measurement of core body temperature through implanted intra-abdominal data loggers is a relatively simple and feasible technique that provides far greater insight into an individual’s thermal balance with its environment.

When alternative methods are used to measure an animal’s surface or peripheral tissue temperature, one should acknowledge that these data do not necessarily reflect the thermal status of an animal’s core in a given environment. For subcutaneous data logging methods, in particular, the limited scientific value afforded by these methods should be weighed carefully against the ethical considerations surrounding the required animal capture and intervention. When it is not possible to measure body temperature within the animal’s core, we suggest that alternative, non-invasive physiological measures are used to provide information on a primate’s energy balance, hormone levels, or metabolic activity in response to environmental variability (e.g., Cristóbal-Azkarate, Maréchal, Semple, Majolo, & MacLarnon, 2016; Behringer & Deschner, 2017; Thompson, 2017; Thompson et al., 2017b). Biophysical models (e.g., Niche Mapper™: Porter & Mitchell, 2006) that use principles of heat and mass transfer, coupled with information on an animal’s morphology, behavior, and microclimate, may also be used to predict an animal’s energetic requirements as a function of environmental conditions (e.g., Moyer-Horner, Mathewson, Jones, Kearney, & Porter, 2015; Natori & Porter, 2007; Zhang, Mathewson, Zhang, Porter, & Ran, 2018; Long et al., 2014; Mathewson & Porter, 2013; Briscoe, Kearney, Taylor, & Brendan, 2016; Mathewson et al., 2018), including in vervet monkeys (Mathewson, 2018). Ultimately, an integrative approach to understanding heat transfer and physiological plasticity can complement accurate measures of core body temperature, to provide greater insight into how primates respond to environmental stress.

## ACKNOWLEDGEMENTS

We thank the Tompkins family for permission to work on the Samara Private Game Reserve. We are grateful to Sr. Mary-Ann Costello, and Drs. Jessica Briner and Anna Haw for veterinary assistance in the animal capture, sedation, and data-logging surgery procedures. Abigail Chiesa helped analyse the thermal images using FLIR software. This research was funded by Faculty research grants from the University of the Witwatersrand, N.S.E.R.C. awards to S.P.H. and L.B., South African National Research Foundation grants to A.F., R.S.H., and S.P.H., and a Carnegie grant to A.F.

## Data sharing

The data and analysis code that support the findings of this study are openly available on figshare at https://doi.org/10.6084/m9.figshare. ####### [to be published at time of manuscript publication].

